# Bite size of *Caenorhabditis elegans* regulates feeding, satiety and development on yeast diet

**DOI:** 10.1101/473256

**Authors:** Atreyee De, Amit K Sahu, Varsha Singh

## Abstract

In the wild, the soil dwelling *nematode Caenorhabditis elegans* primarily feeds on microbes which are abundant in rotting vegetation. Recent published studies showthat several Gram-positive and Gram-negative bacterial populations predominantly constitutes the *C. elegans* gut microbiome but surprisingly lack any yeast species. Here, we show that *C. elegans* display low satiety on yeast diet of *Cryptococcus neoformans, C. laurentii* or *S. cerevisiae*. We found that average size of budding yeast cells is much larger than *E. coli* cells. Yeast cells also cause pharyngeal obstruction, diminished feeding, and lower level of neutral lipids in adult *C. elegans*. Using scanning electron microscopy, we show that the mouth size of *C. elegans* larvae is smaller than average yeast cell. The larvae have no detectable yeast in their alimentary canal and they undergo delayed development on yeast diet. We propose that microbial cell size or bite size could be one of the crucial factors in regulation of feeding in *C. elegans*.

**IMPORTANCE:** The microbiome in *C. elegans* gut is composed of diverse genera of bacteria but it lacks yeast and other fungi. In this study, we provide evidence that yeast cell size is bigger than the mouth size of *C. elegans* larvae. We propose that “bigger than the bite’’ size of yeast cells is one possible reason for low satiety on yeast diet, reduced feeding, lower stored lipids and delayed development. The bite size threshold imposed by *C. elegans* mouth can, partly, explain absence of yeast in *C. elegans* native gut microbiome and “bite size’’ can be studied further as a determinant of microbiome diversity in other animals.

## *Caenorhabditis elegans* displays low satiety on yeast diet

Recent studies report presence of different genera of bacteria from phyla Proteobacteria, Bacteriodetes, Firmicutes and Actinobacteria in *C. elegans* native microbiome but lack of yeast species in spite of presence of fungi in the niches that nematodes were isolated from (1, 2). We hypothesized that *C. elegans* prefers bacteria over yeast and does not feel satiated when only yeast is present. However, *C. elegans* has been shown to have yeast cells in the gut when given *Saccharomyces cerevisiae* (3) or *Cryptococcus neoformans* (4) as diet under laboratory conditions. We found that *C. elegans* adults fed GFP expressing *C. neoformans H99α*::GFP had yeast cells in the gut (Fig. 1A), indicating that adult animals were indeed capable of feeding on yeast cells. We could also detect 700-1000 live yeast/ animal by enumeration of colony forming units (CFU) of yeast (Fig. 1B). To test satiety of *C. elegans,* we studied exploration behaviour of animals on *E. coli* and *C. neoformans* lawns. On food, *C. elegans* displays two distinct foraging behaviours, roaming and dwelling (illustrated in Fig. 1C, shown in Fig. 1D). Dwelling in a localized area indicates food preference, whereas roaming indicates search for better diet and lower satiety (5, 6). Nematode’s roaming response increases with time on laboratory diet of *E. coli* (Fig. 1E). We found that the animals on *C. neoformans* lawn roamed much more than animals on *E. coli* lawn, at 2 hours and 4 hours (Fig. 1F). This indicated that animals may not perceive *C. neoformans* as high-quality food. To find out if virulence of *C neoformans* is a trigger for roaming behaviour in *C. elegans*, we performed exploratory assay with a pathogenic bacterium *Salmonella typhimurium*. However, we found that *C. elegans* exploration on pathogenic bacterium *S. typhimurium* was only slightly higher than on *E. coli* (Fig. 1G). This indicated that a factor other than virulence may be responsible for increased roaming of *C. elegans* on the yeast diet.

**FIG 1.**
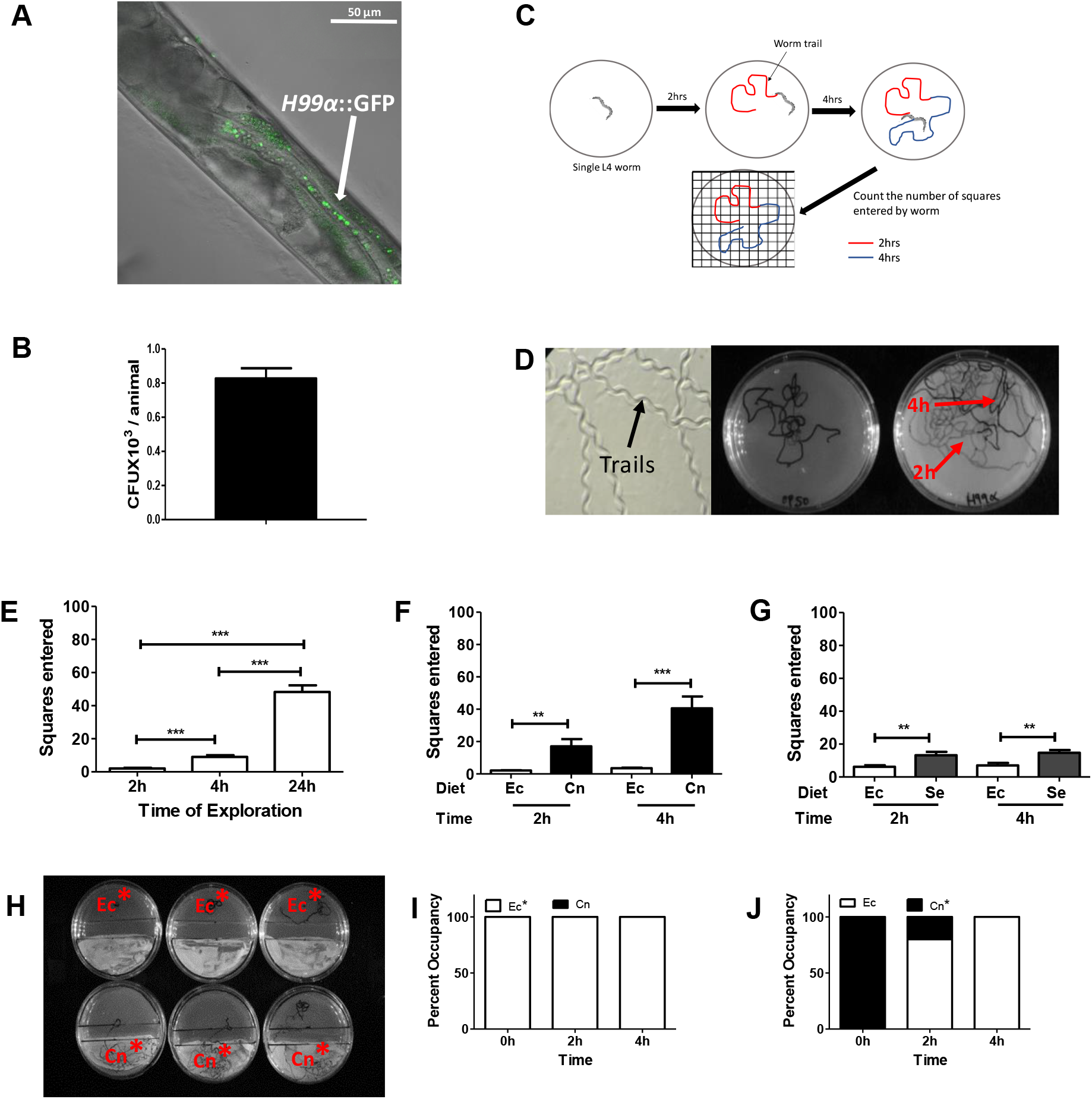
Roaming behaviour and preference of *C. elegans* on budding yeast *C. neoformans*. (A) *C. neoformans H99α*::GFP in the intestinal lumen of adult animals after 24 hours of feeding on the yeast. (B) CFU of *H99α*::GFP recovered from *C. elegans* adult after 24 hours of feeding on yeast. (C) Schematic representation of *C. elegans* exploratory assay quantified using a grid pattern (see methods). (D) Representative trail of adult *C. elegans* on *E. coli*. (E) Quantification of *C. elegans* exploration on monoxenic lawn of *E. coli*. (F) Quantification of *C. elegans* exploration on *E. coli* and *C. neoformans.* (G) Quantification of *C. elegans* exploration on *E. coli* and *S. typhimurium.* (H) Representation of half lawn preference assay for *C. elegans*. (I) Half lawn occupancy of animals released on *E. coli* (Ec*) at 2 and 4 hours, and (J) Half lawn occupancy of animals released on *C. neoformans* (Cn*) at 2 and 4 hours. Unpaired *t* test was used for analysis of significance (ns, P>0.05; *, P<0.05, **, P<0.001; ***, P<0.0001).

To understand if increased roaming on *C. neoformans* lawn was an indicator of low preference of *C. elegans* for yeast, we set up a *C. elegans* preference assay, with half lawns of *C. neoformans* and *E. coli* as choices (Fig. 1H). We released single L4 animal on either *E. coli* (Ec*) or *C. neoformans* (Cn*) half lawn and monitored their choice at 2 hours and 4 hours. We found that all animals released on *E. coli* (Ec*) stayed on *E. coli* half lawn up to 4 hours (Fig. 1I), whereas 8 out of 10 animals released on *C. neoformans* (Cn*) half lawn had moved to *E. coli* half lawn within 2 hours. By 4 hours all animals released on *C. neoformans* lawn (Cn*) had moved to *E. coli* (Fig. 1J) and engaged in dwelling behaviour there (data not shown). Thus, *C. elegans* displayed a clear preference towards *E. coli* over *C. neoformans*. Taken together, our experiments show that *C. elegans* displays low satiety on *C. neoformans* as sole diet as well as low preference over *E. coli*, in a food choice experiment.

## Yeast diet restricts feeding and larval development in *C. elegans*

Low preference of *C. elegans* for *C. neoformans* diet compared to *E. coli* diet could be an indicator of low quality of food as described before for other diets (5). Low quality and quantity of diet can affect larval development due to possible nutrient restriction (6,7). We tested the effect of yeast diet on *C. elegans* larvae. We allowed L1 larvae to feed on a monoxenic lawn of *C. neoformans* or of *E. coli* and monitored development by measuring body length over a period of 72 hours at 25°C (Fig. 2A). Larvae increase in body length as indicated on the right Y axes in Fig. 2B. We observed that larvae fed *C. neoformans* were significantly smaller compared to larvae fed *E. coli* at all time points (Fig. 2A). L1 larvae fed on *E. coli* for 72 hrs had an average body length of 955 µm whereas *C. neoformans* fed larvae did not grow beyond 700 µm in length with an average body length of just 561 µm (Fig. 2B). L2, L3 and L4 larvae provided *C. neoformans* only diet also showed delayed development compared to *E. coli* fed larvae (Fig. S1A, B and C). This indicated that *C. neoformans* diet induced slower growth in *C. elegans* larvae, with the effect becoming less severe with successive larval stage.

**FIG 2.**
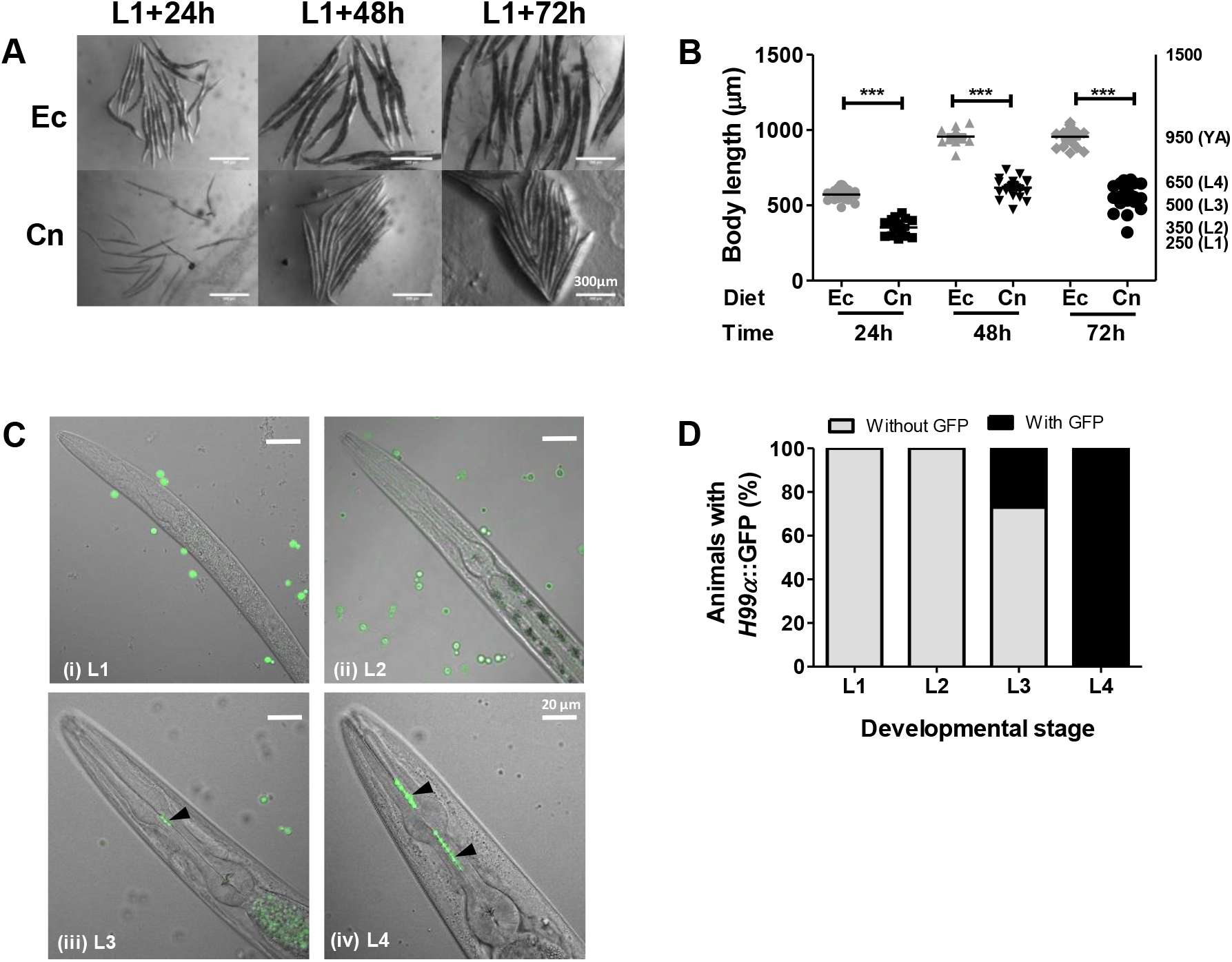
Yeast diet restricts feeding and larval development in *C. elegans.* (A) imaging of L1 larvae fed *E. coli* or *C. neoformans* at 24, 48 and 72 hours. (B) Kinetics of growth represented by body length measurement of L1 larvae fed *E. coli* or *C. neoformans* for 24, 48 and 72 hours. (C) Imaging of L1, L2, L3 and L4 larvae fed *H99α*::GFP for 8 hours. Pharyngeal obstruction or String of yeast cells in the pharyngeal lumen in L3 and L4 larvae are indicated with arrowheads. (D) Quantification of larvae showing presence of *H99α*::GFP in pharyngeal lumen after 8 hours of feeding. 10-20 animals were used for body length measurement at every time point on each diet in panel B. 10-15 animals were used for quantification of pharyngeal yeast shown in panel D. Unpaired *t* test was used for analysis of significance (ns, P>0.05; *, P<0.05, **, P<0.001; ***, P<0.0001).

We noted that diet induced effect on development was not discrete but heterogeneous spanning body sizes from L2 to L4 larval stage (Fig. 2A and B). This indicated that there was no developmental block at a specific point, rather a delay (Fig. 2A). This may result from poor feeding and caloric restriction in the population. To check for *C. elegans* ability to feed on yeast, we allowed the adult and larvae to feed on *H99α*::GFP. We observed that all L4 larvae had ingested yeast in their intestinal lumen within 8 hours of feeding. In addition, we observed retention of strings of yeast cells in pharyngeal lumen in all L4 larvae fed *H99α*::GFP (Fig. 2C). We termed this a pharyngeal obstruction event, not described for *C. elegans* on other diets. We did not detect yeast cells in L1 and L2 larvae fed *H99α*::GFP, either in the gut or in the pharynx (Fig. 2C i –ii and Fig. 2D), however some L3 larvae had *H99α*::GFP (Fig. 2C iii). This suggested that *C. elegans* larvae feed on *C. neoformans* poorly which may lead to nutrient restriction and delayed progression through larval development. Starvation or nutrient restriction in *C. elegans* leads to utilization of neutral lipids stored in lipid droplet, detected by neutral lipid stains such as Oil Red O stain (8). Poor feeding on yeast diet should lead to reduced lipid stores in yeast fed animals compared to well fed animals. Indeed, we found that, yeast fed animals had 40% lower oil red O stained lipids than in *E. coli* fed animals (Fig. S2A and B). Thus, we found that *C. elegans* larvae fed poorly on yeast diet and underwent delayed development. Lower neutral lipid stores in adults indicate that they also derive less nutrition from yeast diet possibly due to reduced feeding. Pharyngeal obstruction caused by yeast cells in bigger animals suggested that yeast cell size may be a contributing factor for reduced feeding.

## Multiple yeast species regulates larval development and satiety in *C. elegans*

Pharyngeal accumulation of *C. neoformans* cells in *C. elegans* L4 larvae indicated that yeast size may impact feeding. Indeed, the reported size of ovoid yeast cell is much bigger than *E. coli*, a Gram negative bacillus. We found that the length of *E. coli* cells measured 2 ± 0.03 µm (Fig. 3A and B) as reported previously (9). The major axis of *C. neoformans cells*, on the other hand, was 5.4 ± 0.2 µm (Fig. 3B) while the minor axis was 4.5 ± 0.1 µm (Fig. 3C). We hypothesized that the big cells size of *C. neoformans* cells prevents *C. elegans* larvae from feeding adequately prompting them to engage in roaming behaviour. If this was true, then other “yeast only’’ diet should also cause developmental delay in *C. elegans* larvae and should induce roaming behaviour in adults. We imaged and measured the cells size of common budding yeast *S. cerevisiae* (Sc) (2) and non-pathogenic yeast *Cryptococcus laurentii* (Cl) (3) (Fig. 3A). The major axis of *S. cerevisiae*, *C. laurentii* yeast cells measured to be 5.8 ± 0.2 µm and 5.4 ± 0.1 µm respectively and were significantly larger than *E. coli* cells (Fig. 3B). The minor axes were 3.6 ± 0.1 µm for *S. cerevisiae* and 4.1 ± 0.1 µm for *C. laurentii* cells (Fig. 3C). *S. cerevisiae* and *C. laurentii* had dimensions comparable to *C. neoformans* cells. Next, we tested if these yeast had any effect on larva developmental in *C. elegans*. Indeed, we found larvae to be lagging in body length when fed *C. laurentii* or *S. cerevisiae* compared to *E. coli* fed control animals (Fig. 3D and E). This indicated that yeast cells cause developmental delay in *C. elegans* larvae. We performed a *C. elegans* survival assay and found that *C. neoformans* (TD_50_ ~70 hours) was more pathogenic than *S. cerevisiae* (TD_50_ 108 hours) while *C. laurentii* was relatively non pathogenic (TD_50_ 134 hours) (Fig. S3A and B). This indicated that three yeast species cause developmental delay in *C. elegans* larvae in a manner not dependent on pathogenicity of yeast.

**FIG 3.**
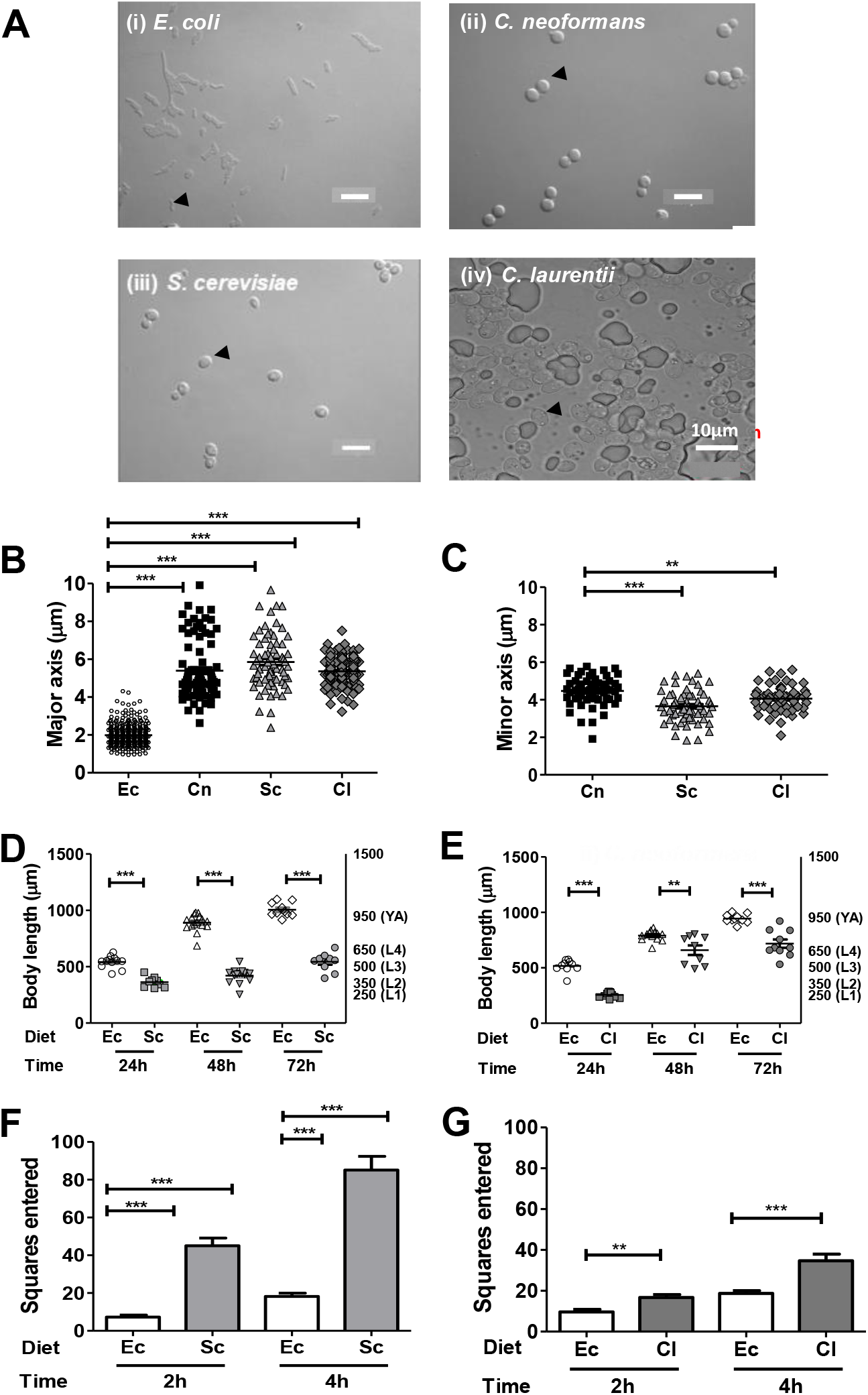
Budding yeast regulate exploratory behaviour and larva development in *C. elegans.* (A) DIC images of *E. coli, C. neoformans*, *C. laurentii* and *S. cerevisiae* cells. (B) Measurement of length or major axes of *E. coli* and yeast species. (C) Measurement of minor axes of yeast species. Kinetics of growth represented by body length measurement of L1 larvae fed (D) *E. coli* or *S. cerevisiae,* (E) *E. coli* or *C. laurentii,* for 24, 48 and 72 hours. Quantification of *C. elegans* exploration on (F) *E. coli* versus *S. cerevisiae* lawns, and (G) *E. coli* versus *C. laurentii* lawn. 70 to 300 cells were used for measurement of major and minor axes in panels B and C. 10-12 animals were used for larval growth measurement in panels C and D. 8-12 animals were used for analysis of exploratory behaviour in panels F and G. Unpaired *t* test was used for analysis of significance (ns, P>0.05; *, P<0.05, **, P<0.001; ***, P<0.0001).

We tested if all yeast causes low satiety in *C. elegans*. In exploratory assays, *C. elegans* showed increased roaming and exploration on *S. cerevisiae* as well as on *C. laurentii,* compared to *E. coli* (Fig. 3G and H; Fig. S4A and B). Roaming behaviour on *C. laurentii* was significantly higher than on *E. coli*, but lower than on *S. cerevisiae* and *C. neoformans* suggesting that roaming behaviour in *C. elegans* may be induced by many attributes of yeast cells. Taken together, our experiments indicate that *C. elegans* is not satiated on yeast, all much larger than *E. coli*, leading to developmental delay in larvae, and enhanced roaming behaviour in adults.

## Inter labial opening in *C. elegans* larvae is smaller than the size of yeast cells

As yeast induced developmental delay was more pronounced in younger larvae, we reasoned that smaller size of mouth in larvae might be the reason for their inability to ingest big yeast cells. *C. elegans* mouth is composed of six labia or lips. Although scanning electron micrographs of adult’s mouth exist (10) there is no study of mouth size through development. Therefore, we studied labia arrangement and mouth size in all larval stages, by scanning electron microscopy. We observed that 6 labia arrangement was present in all stages (marked in L4 larva in Fig. 4A). We also measured the mouth of L1 larva, L2 larva, L3 larva and L4/young adult (Fig. 4B, i-iv) as inter labial distance between two opposing labia. As shown in Fig. 4C, the inter labial distance increased with developmental progression. The mouth size was 1.2 ± 0.03 µm in L1 larvae, 1.6 ± 0.1 µm in L2 larvae, 2.3 ± 0.1 µm in L3 larvae, and 4.1 ± 0.1 µm in young adults. As a control, we also measured the diameter of the animals’ head at the third pharyngeal furrow (indicated in Fig. 4A) which also showed a similar trend of increase through larval stages as did the mouth size (Fig. 4D). Mouth size of 1.2 µm in L1 larvae, in our SEM analysis, is in agreement with previous report which indicated that L1 larvae of *C. elegans* cannot ingest polystyrene beads >1 µm (10). Our data shows that size of mouth in L1, L2 and L3 larvae is much smaller than the yeast cell size of 3.6-6 µm and likely presents a hurdle to ingestion.

**FIG 4.**
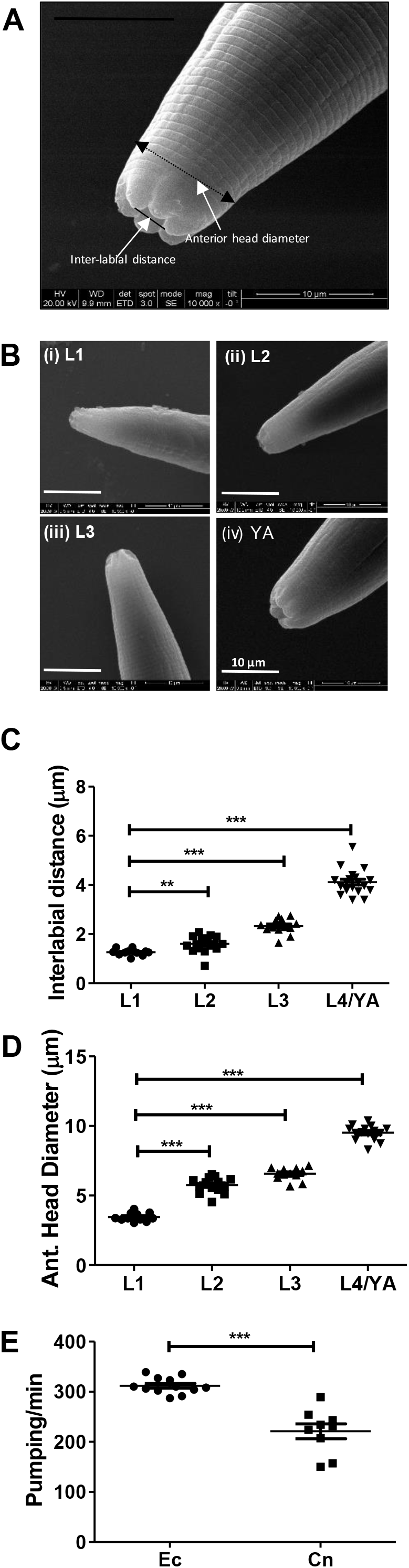
Inter labial opening in *C. elegans* larvae. (A) Scanning electron microscopy (SEM) image of mouth of *C. elegans* L4 larva. Inter labial distance and anterior head diameter are indicated. (B) SEM images of mouth of L1, L2, L3 larvae and young adult (YA). (C) Quantification of mouth size as inter labial distance across developmental. (D) Quantification of anterior head diameter across developmental stages. (E) Pharyngeal pumping in adult animals on *E. coli* versus *C. neoformans* after 8 hours of exposure. 12-20 animals were used for quantification of mouth size and anterior head diameter. 10-15 animals were used for analysis of pumping rates quantified in panel E. Unpaired *t* test was used for analysis of significance in panels C, D and E (ns, P>0.05; *, P<0.05, **, P<0.001; ***, P<0.0001).

If *C. elegans* bite size presents a minimum threshold for ingestion, the nematodes might engage in roaming behaviour on yeast lawn, possibly to find smaller yeast cells for easy ingestion. Therefore, we measured the diameter of yeast cells present in pharyngeal lumen of L4 larvae and found that size of ingested yeast (3.1 ± 0.1 µm) was indeed significantly smaller than average yeast cell size (>4 µm) (Fig. S5). This supported the idea that nematodes feed on smaller yeast cells in a lawn although the mode of sensing of smaller cells by *C. elegans* is not known. We reasoned that nematode’s inability to ingest larger yeast and pharyngeal obstruction should lower pharyngeal pumping motion, necessary for feeding and ingestion (12). We observed that pumping rate in *C. elegans* adults decreased by 30% while feeding on *C. neoformans* compared to *E. coli* pointing to difficulty faced by animals in ingestion of yeast cells (Fig. 4E). We looked for additional evidence to determine if *C. elegans* has a bite size preference in its natural habitat. By literature survey (Fig. S6), we collated reported cell sizes for most genera of bacteria present in *C. elegans* gut microbiota (2). We found that majority of bacteria had average width/minor axis below 1 µm (Fig. S7). This analysis suggested that even in its natural habitat, with a wide diversity of diet choices, *C. elegans* might have preference for smaller microbes.

In summary, our study shows that the mouth size of *C. elegans* increases with development and positively correlates with its ability to ingest yeast cells bigger than 4 µm. Younger larvae, with smaller mouth size, have undetectable yeast in their gut or pharynx and show delayed development pointing to difficulty of young animals in the ingestion of yeast cells. Adult *C. elegans* also shows lower satiety and reduced neutral lipid stores on yeast diet. Thus, we propose that ingestibility of microbes is likely to be one of the criteria for determining preference of *C. elegans* for microbes. Larger mouth or bite size needed to ingest yeast cells may also explain lack of yeast in the native microbiome of this nematode. Study of diet preference of other soil dwelling nematodes which feed on microbes would be useful in substantiating bite size a determinant of microbiome composition.

## METHODS

### *C. elegans* and microbial strains

*C. elegans* N2 Bristol strain was obtained from Caenorhabditis Genetics Centre. The animals were maintained on *E. coli* (OP50) seeded nematode growth media (NGM) plates at 20°C. *Salmonella typhimurium* SL1344 was grown overnight in LB broth and plated on NGM agar. *Cryptococcus neoformans* (H99α, *H99α*::GFP), *C. laurentii* (LS28) and *S. cerevisiae* were grown in yeast peptone dextrose (YPD) broth and plated on Brain heart infusion (BHI) agar.

### Confocal Imaging

*C. elegans* larvae and adults were fed on *H99α*::GFP (RBB1) for 8 hours or 24 hours and imaged with Zeiss LSM 880 Airyscan to visualize fluorescent yeast cells.

### Yeast colonisation of *C. elegans* intestine

*C. elegans* adults were fed on GFP tagged *C. neoformans*. Animals washed 3 times with M9 buffer were mechanically disrupted using a sterile plastic pestle. Colony forming units of yeast were enumerated on brain heart infusion (BHI) agar plates containing gentamycin (50ug/ml) and expressed as number of yeast per animal.

### Exploration assay

Single L4 animals were released on a 60mm NGM agar uniformly seeded with yeast or bacterial strains. Trails left by the animal were traced with color markers. The traces were superimposed on a grid containing 11*11 squares, and the number of squares traversed by the worm tracks were manually counted, adapted from (13). Traces on all microbial strains were compared to traces on control *E. coli* OP50 strain. A minimum of 10 animals were tested per microbe.

### Preference assay

On a 60mm Nematode growth media (NGM) plate, half lawns of *C. neoformans* and *E. coli* were made, with a clearance of 5mm between two halves. Single L4 larvae were either released on *E. coli* lawn or on *C. neoformans* lawn and monitored for change of feeding preference at 2 and 4 hours.

### Larva development assay

Synchronized population of L1, L2, L3, and L4 larvae were fed on *C. neoformans*, *C. laurentii, S. cerevisiae* at 25 °C. Every 24 hours, 10-15 animals were imaged by bright field microscopy at 35X magnification using Zeiss SteREO Discovery V12 microscope. Imaging was done for three consecutive days and the body length of animals was measured using Image J software.

### Microbe size measurement

Log phase culture of yeast and bacterial samples were taken and imaged using Leica DMI8 microscope. Cell sizes were measured using Leica LAS X imaging software. *C. laurentii* cells were imaged using Zeiss LSM80 Aryscan and cell sizes were measured using Zeiss ZEN software.

### Survival Assay

For survival assay on yeast strains, *C. neoformans*, *C. laurentii*, *S. cerevisiae,* were grown in YPD broth overnight at room temperature. Yeast culture was spotted on BHI gentamycin (50μg/ml) agar plates and incubated at 28°C for 12-14 hours. Young adults of *C. elegans* were transferred to yeast lawns (50-60 worms/plate). Animals were transferred to fresh yeast plates every day and scored as live or dead at regular intervals of time throughout the course of the assay.

### Oil red O Staining for neutral lipid droplets

Solution of Oil Red O (SIGMA ALDRICH) was prepared in isopropanol (0.5g/100ml) and diluted to 60% in water before use. Synchronized L4 animals were allowed to feed on *E. coli* and *C. neoformans* lawns for 8 hours. *C. elegans* adults were fixed and permeabilized using MRWB buffer for 1 hour at room temperature (7). The animals were stained with 60% ORO at room temperature. Stained animals were mounted on slides and imaged using Olympus IX81 bright field microscope. ORO staining was quantified using Image J software.

### Scanning electron microscopy

Animals were washed three times in PBS, and then incubated in 1 ml fixation buffer (0.2mM HEPES, 5% glutaraldehyde, 4% formaldehyde) (14) for 18 hours at room temperature. The fixed animals were washed, twice with PBS and twice with water. Fixed animals were subjected to serial ethanol dehydration (10%/30min, 30%/60min, 50%/30min, 70%/60min, 80%/30min, 90%/30min and 100%/60min). Samples were mounted on a 10% hydrochloric acid coated coverslip followed by gold coating to visualise under ESEMQuanta scanning electron microscope. Anterior head diameter and inter labial distance were measured using Image J software.

### Pharyngeal pumping

Pharyngeal pumping was measured by direct counting under Nikon SMZ745 stereomicroscope for 30 seconds. At least 10 animals were monitored for each assay.

## ACKNOWLEDGEMENT

We thank Joseph Heitman, Duke University Medical Center for providing us *C. laurentii* and *C. neoformans H99α*::GFP strains. We thank Kaustav Sanyal, JNCASR, Bangalore for *C. neoformans* H99*α* and Purusharth Rajyaguru, Indian Institute of Science for providing *S. cerevisiae* strain. *C. elegans* strains were provided by the CGC, which is funded by NIH Office of Research Infrastructure Programs (P40 OD010440). Atreyee De is supported by scholarship from University Grants Commission of the Government of India. This work was partially supported by the Welcome Trust/DBT India Alliance Intermediate Fellowship (Grant no. IA/I/13/1/500919 2013-2018) awarded to Varsha Singh.

